# Consequences of alternative stable states for short-term model-based control of cyanobacterial blooms

**DOI:** 10.1101/2023.12.15.571830

**Authors:** Bas Jacobs, George van Voorn, Peter van Heijster, Geerten M. Hengeveld

## Abstract

Cyanobacteria can form dense blooms in eutrophic lakes that can be toxic to humans and other animals and harmful to the ecosystem’s functioning. While better nutrient management is generally considered the long-term solution to this problem, short-term mitigation efforts (e.g., flushing, algaecides, flocculants) are becoming increasingly necessary to safeguard water quality and the ecosystem services it provides. Here, we explore potential model-based management strategies for these short-term mitigation efforts. We focus on the case where blooms are linked to the existence of alternative stable states, such that, under the same conditions but depending on the past, a lake may be dominated either by cyanobacteria (causing a harmful algal bloom) or by green algae and macrophytes in a clear water state. Changing conditions may cause the favourable clear water state to disappear through a tipping point, causing the lake to switch rapidly to the turbid cyanobacteria state. At the same time, it may take considerable effort to undo this tipping and return to the favourable state. We identify four different strategies for bloom mitigation in this scenario: Doing nothing, reacting to a bloom, resetting the lake at a later point, and preventing the bloom. We explore the different requirements for these strategies along with their associated cost profiles. We also investigate the effect of transition times from one state to another on the efficacy and costs of different strategies.

## 1. Introduction

Cyanobacteria (or ‘blue algae’) are photosynthetic bacteria that can cause major problems in eutrophic lakes due to their ability to form dense blooms (Huisman et al., 2018; Burford et al., 2020). These blooms may cause ecological damage by blocking light from submerged macrophytes (Paerl and Otten, 2013). They also pose a health risk, since cyanobacteria can produce a variety of toxins that are harmful for freshwater lifeforms as well as humans (He et al., 2016). Consequently, blooms cause economic damage by, for instance, incurring extra costs for the purification of drinking water and making the water unsuitable for recreational purposes (Merel et al., 2013). Since blooms appear to thrive in warmer weather, they are expected to become an even larger problem as temperatures increase (Paerl and Huisman, 2008).

The long-term solution to the bloom problem is to reduce the introduction of excess nutrients that created the eutrophic conditions favourable to blooms in the first place (Ibelings et al., 2016). However, this solution is often hard to implement and even if the addition of excess nutrients is stopped, it may still take a long time for a lake to recover due to nutrients stored in the sediment (Fastner et al., 2016; Søndergaard et al., 2013). These issues may make short-term mitigation efforts necessary for the foreseeable future (Lürling and Mucci, 2020). These mitigation efforts may take the form of, for example, flow rate control (Mitrovic et al., 2010; Ibelings et al., 2016) or the addition of cyanocidal compounds such as hydrogen peroxide (Matthijs et al., 2016).

Models of aquatic ecosystems have strong potential for decision support at many different levels. Large descriptive models are often used for long-term scenario analysis, e.g., to predict the effect of climate change or long-term management interventions such as changing the nutrient load (Mooij et al., 2007; Janse and van Liere, 1995; Los and Wijsman, 2007). Models can also be used on shorter time scales, e.g., to provide an early warning for an impending bloom several days in advance (Ibelings et al., 2003; Trolle et al., 2014; Page et al., 2018). An obvious application for such a warning would be to post warning signs for swimmers. Short-term predictions also offer potential for the timing of mitigation interventions.

The timing of interventions is especially important when alternative stable states exist, i.e., depending on the starting point, the system can end up in both a clear state dominated by green algae and a turbid state dominated by blue algae (i.e., a bloom). When two such alternative stable states exist (i.e., bistability, see Box 1 for a terminology overview), a small change in environmental conditions and management interventions may remove one of the stable states, creating a tipping point (i.e., bifurcation point). The resulting sudden switch to the other stable state can have large consequences that are difficult to reverse (e.g., hysteresis). Alternative stable states have been found both for small abstract models for cyanobacterial blooms (Scheffer et al., 1997a; Gragnani et al., 1999; Scheffer and Rinaldi, 2000) and large descriptive ones like PCLake (Janse et al., 2010). Experimental evidence for the existence of alternative stable states has been found in many ecological systems (Schröder et al., 2005), though their existence in real freshwater ecosystems remains difficult to prove for individual lakes (Capon et al., 2015). However, sudden transitions between a clear state dominated by green algae and a turbid state dominated by blue algae are observed in lakes, and the early warning indicators based on the concept of ‘critical slowing down’ near bifurcation points (van Nes and Scheffer, 2007; Carpenter et al., 2009; Veraart et al., 2012) appear to work in real ecosystems (Pace et al., 2017; Wilkinson et al., 2018).

### Box 1

**Terminology**

- *Cyanobacteria / blue algae*: Photosynthetic bacteria that can cause harmful blooms, particularly when temperatures are high and nutrients are plenty. Blue algae is a commonly used name for cyanobacteria, even though they are technically not algae.
- *Alternative stable states*: When there are alternative stable steady states, the same environmental conditions can lead to different outcomes for different starting values. For example, starting with a high density of blue algae may lead to a lake dominated by blue algae, while starting with a low density of blue algae leads to a lake dominated by green algae, all other things being equal.
- *Bistability*: Having two alternative stable steady states.
- *Bifurcation*: Point in parameter space where there is a change in the number or quality (e.g., stable or unstable) of the steady states of a system.
- *Tipping point*: Point in parameter space where the system suddenly switches from one state to another as a result of crossing a bifurcation point. For example, a lake may suddenly lose the steady state dominated by green algae and switch to the (remaining) steady state dominated by blue algae.

Here, we explore the consequences of alternative stable states for potential model-based strategies for cyanobacterial bloom management. To this end, we will assume that the lake under consideration shows bistability. We then explore the effectiveness of several potential short-term mitigation efforts, including strategies based on predicting the magnitude and timing of interventions required to either reverse, or prevent, a bloom. We will also explore the cost profiles associated with different strategies and the trade-offs between bloom costs and control costs that these strategies imply. Finally, we will explore how the efficacy and costs of different strategies depend on transition times from one state to another and on the locations of the bifurcation points in a two-dimensional parameter space.

## 2. Methods

### 2.1 The model

Our goal is to explore the effect of having alternative stable states and bifurcations in an ecosystem on potential management strategies. Therefore, the focus is not on any specific model. For demonstration purposes, we will use a simple competition model with green and blue algae that allows for alternative stable states, based on Scheffer et al. (1997a). This model will serve as an example of one of the many aquatic ecosystem models with alternative stable states, in order to show the consequences of their existence for management decisions. We do not aim to analyse this particular model, nor validate it or otherwise compare it to reality. Therefore, we have included all the technical details of the model in Appendix A. For our purposes, the relevant components of the model are that there is an environmental parameter σ that cannot be controlled (e.g., temperature) and a flow rate *f* as a parameter that can be controlled to mitigate blooms (see Table 1 for an overview of important symbols). Changes in these two parameters can move the system between regimes with different stable steady states: a regime with one stable state dominated by green algae, a regime with one stable state dominated by blue algae, and an intermediate regime where both these stable states co-exist, i.e., alternative stable states (Fig. 1). Importantly, we assume that the environmental parameter can vary rapidly (i.e., on a timescale of days) and that the control parameter can be adjusted on a similar timescale. The model also uses a small inflow of both species to prevent complete extinction, allowing states to switch when conditions change.

**Table 1:** Descriptions and units of important symbols.

| Symbol | Description | Units of |
| --- | --- | --- |
| $g$ | Density of green algae | Numbers per volume |
| $b$ | Density of blue algae | Numbers per volume |
| $\sigma$ | Environmental parameter | Example specific (could be temperature, light intensity, etc.) |
| $f$ | Control parameter. Here: a flow rate. | Example specific. Here: volume per time. |
| $\Delta f$ | Control effort, i.e., the change in control rate. | Same as $f$ |
| $t_b$ | Bloom duration | Time |
| $t_c$ | Control duration | Time |
| $x$ | Bloom cost per duration | Cost per time |
| $\xi$ | Control cost per effort per duration | Cost per $\Delta f$ per time |
| $\alpha$ | Control effort of the reset only strategy | Same as $\Delta f$ |
| $\beta$ | Control effort of the reactive strategy | Same as $\Delta f$ |
| $\gamma$ | Control effort of the preventive strategy | Same as $\Delta f$ |

**Figure 1:**
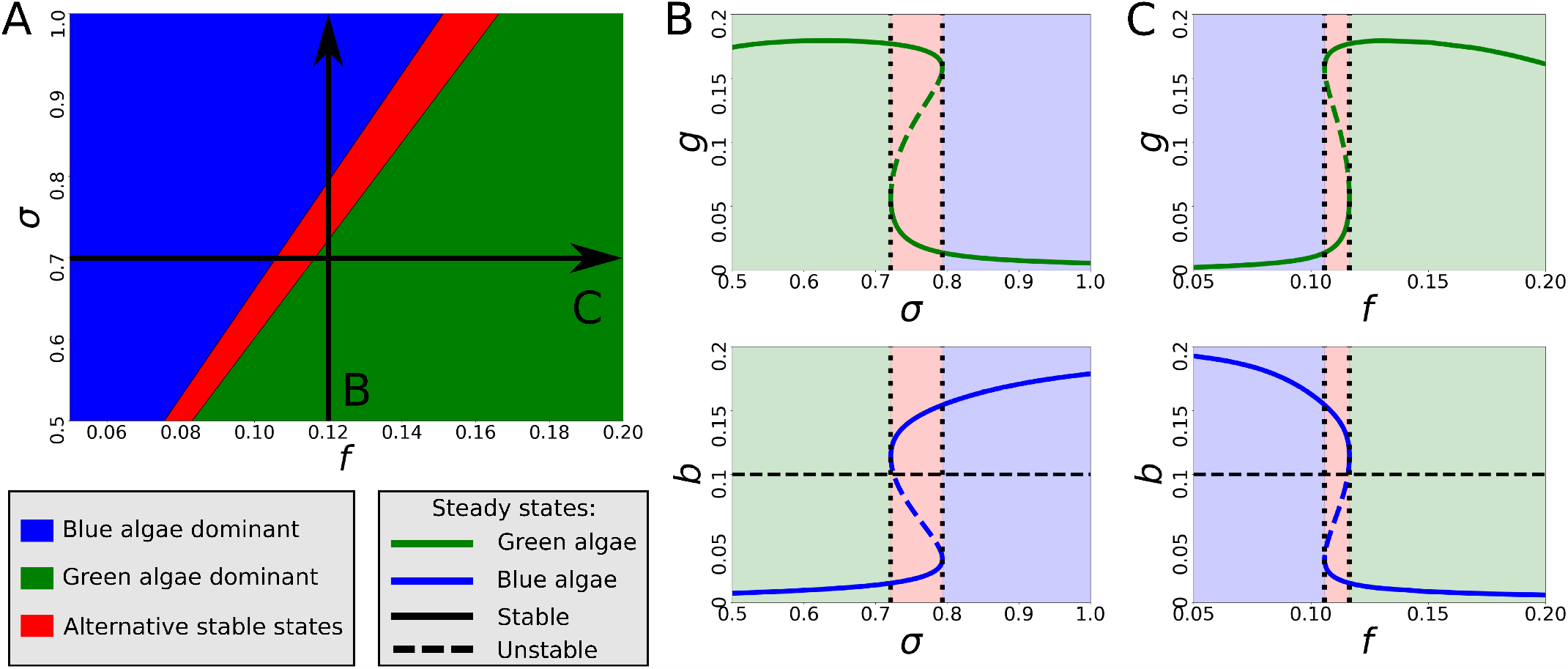
Example bifurcation diagram. (A) Two-parameter bifurcation diagram with control parameter *f* and environmental parameter σ. The green and blue regimes have a single stable steady state dominated by green and blue algae, respectively, and the red regime has two alternative stable states (one dominated by green algae and one by blue algae), where the past determines which state the system is in. The different regimes are separated by fold bifurcations. (B,C) Steady states of green algae (top) and blue algae (bottom) along the sections indicated by arrows in (A). Solid lines indicate stable steady states and dashed lines indicate unstable ones, which exist in the bistable region (red). Background colours correspond to the regimes in (A). Horizontal dashed lines indicate the density of blue algae above which we consider there to be a bloom.

### 2.2 Simulations and bifurcation analysis

Simulations were performed in Python 3. To find and track fold bifurcations through parameter space we used the continuation software AUTO-07p (Doedel et al., 2007).

### 2.3 Bloom scenario and potential management strategies

For lakes that are solidly in a green-dominated or blue-dominated regime, short-term mitigation efforts are either unnecessary or insufficient. For lakes without tipping points, where a gradual transition from one state to the other occurs, the added value of a model is relatively low, since, e.g., intervention timing is far less critical without tipping points. Therefore, we will consider a scenario where we start with a lake in a parameter regime where two alternative stable states exist (one dominated by green algae, the other by blue algae). The lake starts in the (preferred) state dominated by green algae. We then consider a fixed temporary increase in environmental parameter σ that takes the system across a bifurcation point and causes the system to switch to a cyanobacteria-dominated state (a ‘bloom’). Due to the bistability of the system, without further intervention, this leaves the lake in a cyanobacteria-dominated state even after σ has returned to its original value.

We consider four different management strategies for dealing with this scenario: *no intervention, reset only, reactive*, and *preventive*. The *no intervention* strategy involves doing nothing and leaving the lake dominated by blue algae. It does not require any resources or model predictions. The remaining three strategies involve actual interventions, and require a reliable estimation of the actual bifurcation structure of the lake. The *reset only* strategy involves waiting for the environmental parameter to return to its base level and then temporarily increasing control parameter *f* to reach the regime with only a green algae-dominated stable state. The *reactive* strategy involves increasing *f* after the system has switched to a cyanobacteria-dominated state but before σ has returned to its base level. The *preventive* strategy involves pre-emptively increasing *f* to prevent the system from ever crossing the bifurcation point into the regime with only the cyanobacteria-dominated state. This strategy additionally requires a reliable estimation of the oncoming change in σ.

### 2.4 Management strategy costs

We explore the cost profiles of the different management strategies by looking at the required control effort, the time during which this effort must be applied, and the time during which a bloom is present. The control costs and bloom costs could be interpreted as financial costs (e.g., loss of revenue from recreation) but also more broadly (e.g., loss of biodiversity, or aesthetics). For simplicity, we assume that the control cost is directly proportional to the required increase ∆ *f* in control parameter *f* . We also assume that this control cost per unit time ξ and bloom cost per unit time *x* both remain constant over time. We consider a bloom to be present when the density of blue algae exceeds a specific threshold (horizontal dashed lines in Fig. 1B,C).

## 3. Results

### 3.1 Exploration of management strategies

We take a starting point in the regime with two alternative stable states (the red area in Fig. 1 and 2). Without intervention, the increase in environmental parameter σ eventually switches the lake to a cyanobacteria-dominated state (i.e., causes a bloom) that remains even after σ returns to its starting level (Fig. 2A). The *reset only* strategy results in a bloom that lasts until the environmental conditions become favourable again, at which point the temporary increase in control parameter *f* switches the lake back to a state dominated by green algae (Fig. 2B). The *reactive* strategy briefly allows the bloom to appear, followed by an intervention with a larger increase in *f* (compared to the reset only strategy) that switches the lake back to the green state while σ is still elevated (Fig. 2C). This approach requires that the control parameter remains elevated until σ returns to its starting level. The *preventive strategy* requires that the increase in σ can be predicted, but can prevent the bloom entirely by increasing *f* to keep the lake out of the regime with only a blue algae dominated stable state when the environmental parameter σ increases (Fig. 2D). This approach also requires that the control parameter remains elevated until σ returns to its starting level, but the initial control effort required is smaller than for the reactive strategy.

**Figure 2:**
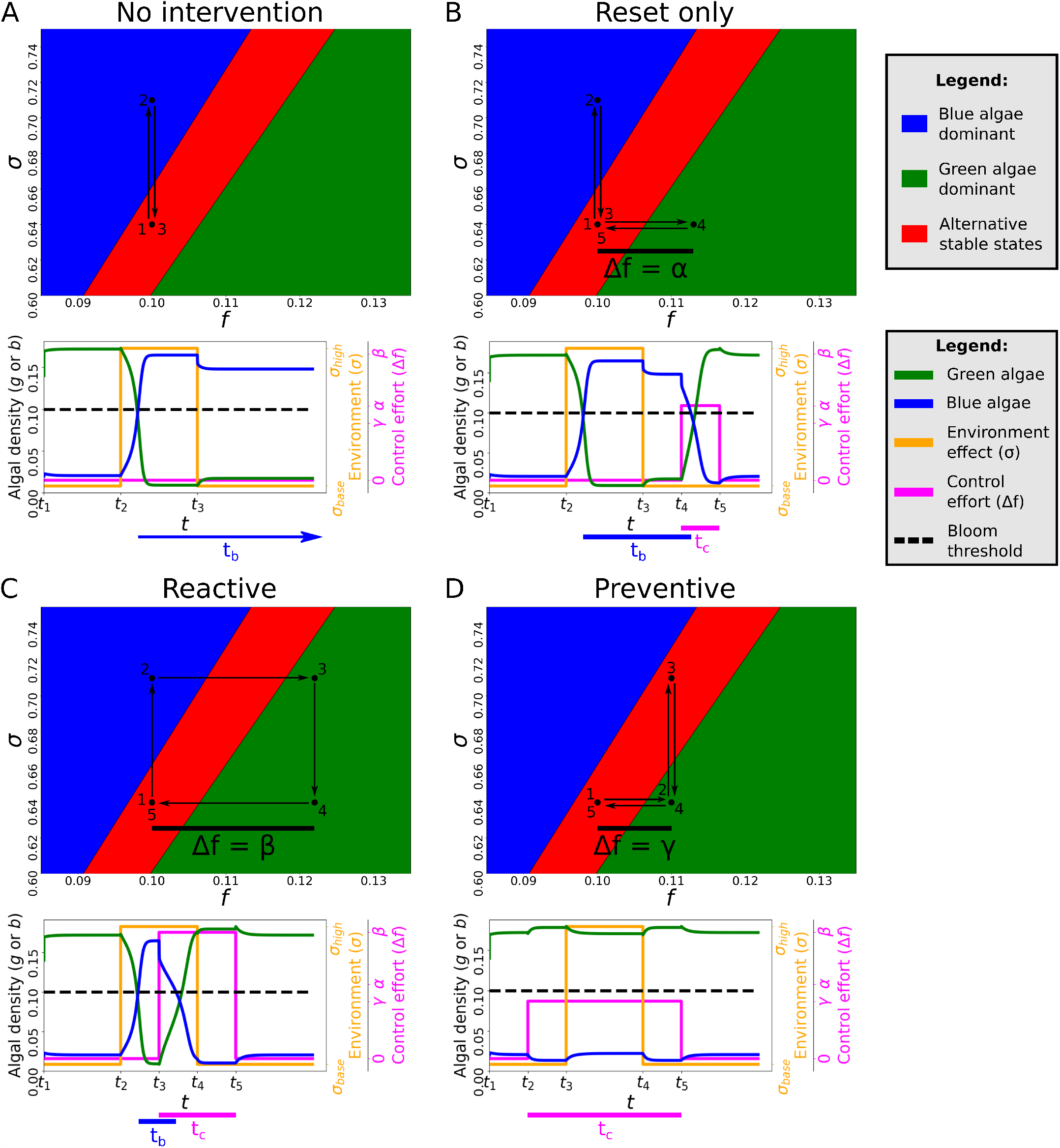
Four potential management strategies for blooms that result from an environmental parameter σ temporarily crossing a bifurcation point. The system is assumed to start in the bistable regime (red region) in a state dominated by green algae. Arrows and numbers in the bifurcation diagrams indicate the order in which the parameters change. The plots below show the transient behaviour of the densities of green algae (*g*) and blue algae (*b*) in response to the changes in the environmental parameter σ (orange) from σ_*base*_ tot σ_*high*_ and back. The control effort (∆ *f*, purple) denotes the increase in *f* from its base level. These increases are labelled α, β, and γ for the reset, reactive, and preventive strategies, respectively. The dashed line indicates the density of blue algae above which we consider there to be a bloom. Time points *t*_1_–*t*_5_ correspond to the numbers in the bifurcation diagram. Time spans *t*_*b*_ and *t*_*c*_ indicate the duration of the bloom and the duration of the control effort, respectively. (A) Without intervention, the change in environmental parameter σ causes a bloom, which persists upon return to the bistable regime. (B) After the system has returned to the bistable regime, it can be switched back to the green state by temporarily increasing control parameter *f* (the flow rate) into the green regime. (C) Reversing the bloom directly after it has occurred requires increasing *f* to reach the green regime and then keeping it there until the environmental parameter has returned to its original value. (D) If the change in the environmental parameter can be predicted, then the bloom can be prevented by increasing *f* before the environmental change to make sure the system never enters the blue regime.

### 3.2 Costs of blooms and management

If we assume that the control cost per unit time is directly proportional to the required control effort ∆ *f*, then the total intervention cost depends on ∆ *f*, the control duration *t*_*c*_, and the control cost per effort per time ξ. The decision maker can make a trade-off between the intervention costs and the costs of the bloom, which includes both financial costs, such as increased purification costs or reduced revenue from recreation, and costs that are harder to express in monetary value, such as nuisances from bad odours or loss of biodiversity. Assuming that a reasonable estimation of this bloom cost can be obtained, the total cost can be calculated from the bloom cost per unit time *x* and the duration *t*_*b*_ of the bloom. The overall total cost is the sum of the cost of the management intervention and the cost of the bloom:

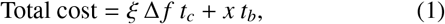

where ∆ *f, t*_*c*_, and *t*_*b*_ are strategy-dependent. In our example, the control effort ∆ *f* is a flow rate expressed units of volume per time, but it could be any quantity with any unit, as ξ is expressed per unit of effort.

The value of control effort ∆ *f* in our example was 0, α, β, or γ, depending on the strategy, resulting in different cost profiles (Fig. 3). The minimum required control effort β of the reactive strategy is always greater than that of the other two strategies, but the effort α for the reset strategy can be greater or smaller than the effort γ for the preventive strategy depending on the structure of the bifurcation diagram. The time *t*_*b*_ during which a bloom exists is largest when not intervening and the smallest (zero) when preventing the bloom altogether. Of all the strategies that include actual intervention, the time *t*_*c*_ during which control effort is required is the shortest when only resetting the system and the longest when preventing the bloom altogether (Fig. 3).

**Figure 3:**
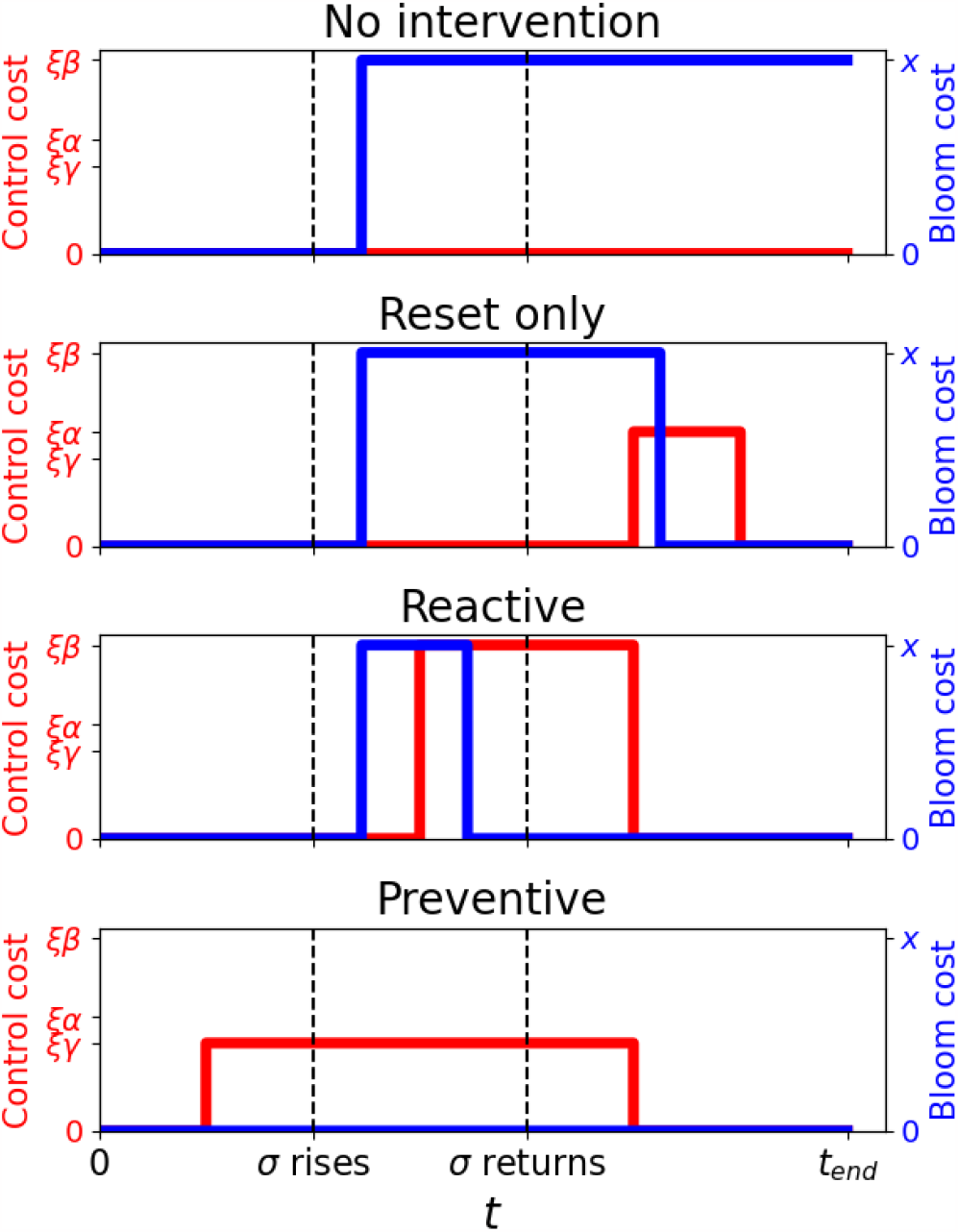
Cost profiles for management costs (red) and bloom costs (blue), associated with the different management strategies shown in Fig. 2. The total costs can be obtained by adding the areas under the bloom and control cost curves. The dashed lines correspond to the points at which environmental parameter σ is increased and reduced, respectively. The cost per time of a bloom (*x*) is assumed to be constant over time, as is the cost per control effort per time *ξ*. The control efforts *α, β*, and *γ* correspond to those in Fig. 2.

### 3.3 Effects of transient dynamics

Near a bifurcation point, transient dynamics are often slow, suggesting that a switch from one state to another can take longer if the bifurcation point is crossed by smaller margin. Therefore, increasing the magnitude of the intervention effort may reduce the required intervention time, as well as the bloom time, particularly when using the reset strategy (compare Fig. 4A and B). Conversely, when trying to reverse a bloom, enough time should be given to allow the switch to the new steady state, otherwise, the reversion may be unsuccessful (Fig. 4C).

**Figure 4:**
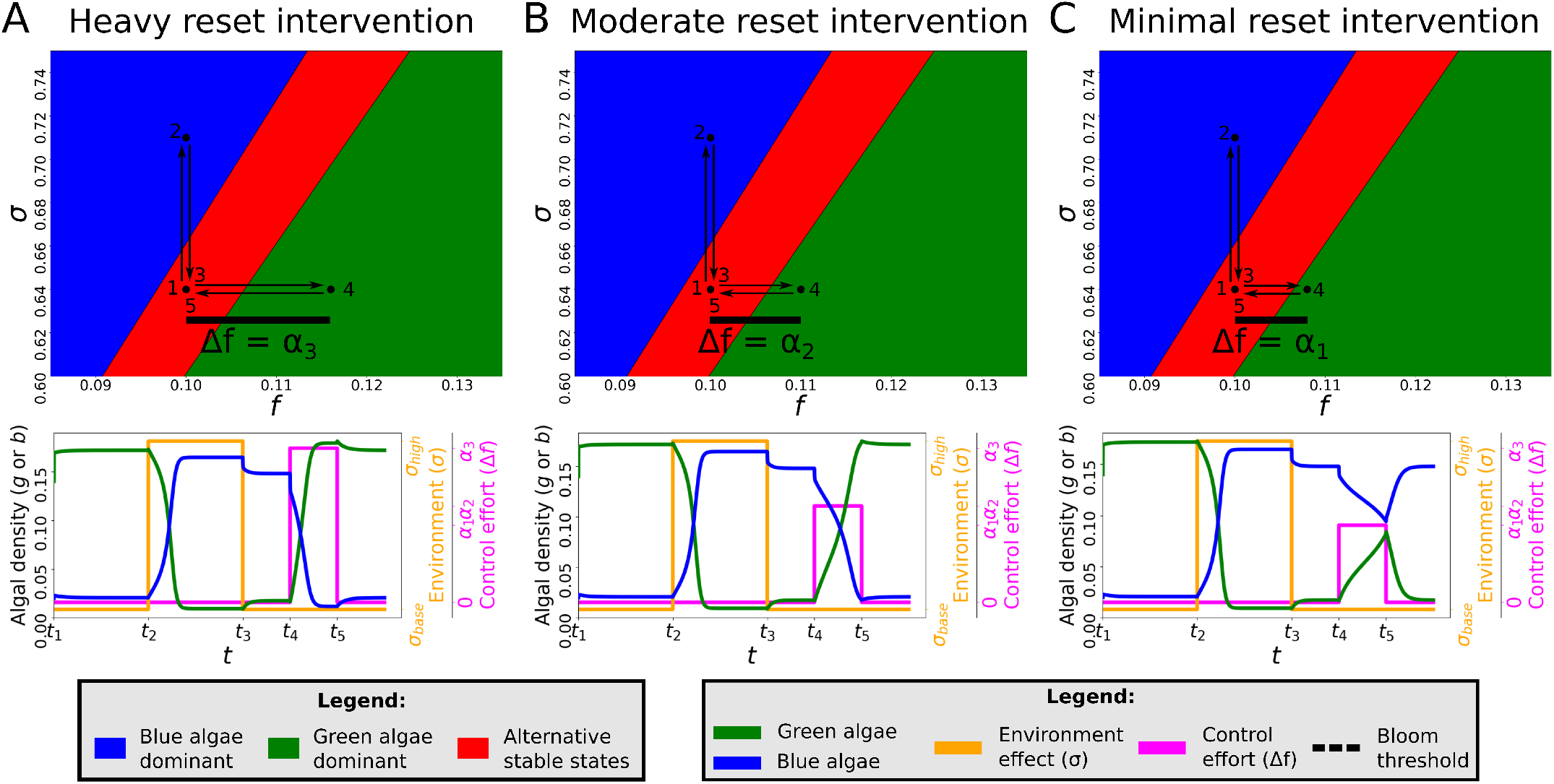
Effect of transient behaviour for different reset intervention intensities. Top panels indicate the three different control efforts ∆ *f* . Bottom panels show the transient behaviour. Meanings of labels and colours are as in Fig. 2. Under heavy intervention (A), the system rapidly reverts to a state dominated by green algae. With a smaller increase in *f* (B), the systems takes longer to revert (as can be seen from the steepness of the green curve). With an even smaller increase (C) the reversion takes even longer and the intervention time is not long enough to make the switch back to the steady state dominated by green algae (i.e., the intervention has been unsuccessful).

### 3.4 Effects of the bifurcation structure

The bifurcation structure of the lake under consideration determines the required control effort for, and feasibility of, the different management strategies. The effect of different bifurcation structures (with straight bifurcation lines for simplicity) is illustrated in Fig. 5. Differences in the steepness of the left bifurcation line change the effort needed to stay in the bistable regime and therefore the cost of the preventive strategy. The steepness of the right line affects the effort needed to move to the regime dominated by green algae after the environmental change and therefore the cost of the reactive strategy. The distance between the bifurcation lines influences the effort needed to move to the green regime regardless of the environmental parameter and therefore affects the costs of both reactive and reset strategies.

**Figure 5:**
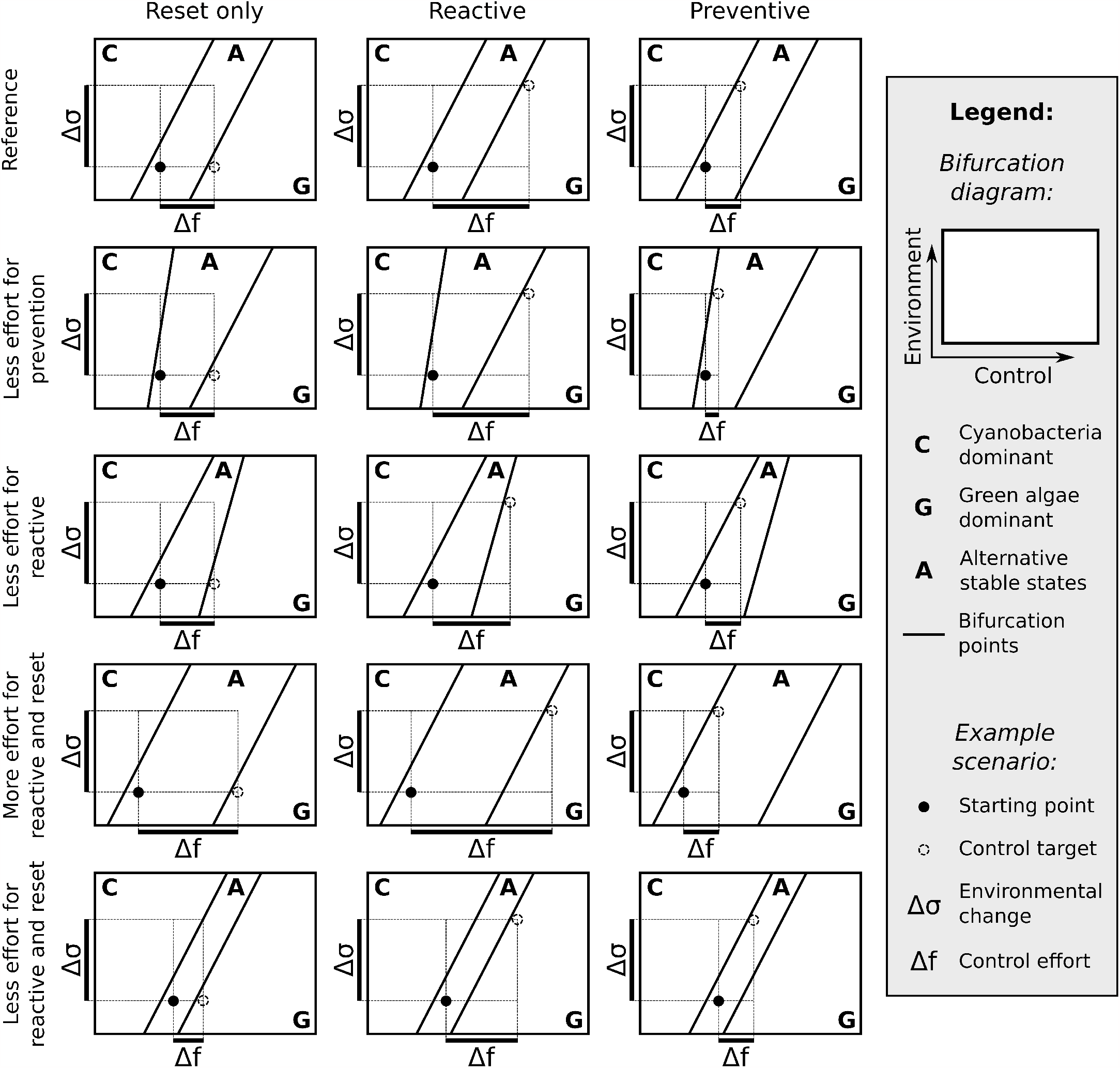
Effect of the bifurcation structure on management strategies. Cartoons show different potential bifurcation structures compared to a reference bifurcation structure similar to that in Fig. 1 (only with perfectly parallel lines), as well as an example management scenario for the reset, reactive, and preventive strategies. The environmental change ∆σ is kept constant, while the control effort ∆ *f* is adjusted to reach the minimal control target for the relevant strategy (open circles). First row: Reference diagram showing the minimal control targets for the three different strategies. Second row: The steepness of the left line impacts the minimal control effort for the preventive strategy. Third row: The steepness of the right line impacts the minimal control effort for the reactive strategy. Fourth and fifth row: The distance between lines impacts the minimal control effort for the reactive and reset strategie

## 4. Discussion

Our exploration revealed several distinct strategies for bloom management in lakes with alternative stable states, as well as some key factors in deciding between these strategies. The optimal choice of strategy depends on a variety of factors including the bifurcation structure of the actual lake under consideration, the costs per time of a bloom and the proposed intervention, and the transient effects that determine bloom duration and the required duration of an intervention. For example, if the costs of a bloom are considered unimportant, then the best strategy will be *no intervention*, as this minimises the intervention costs. Conversely, if blooms are completely unacceptable, the *preventive* strategy will be the best option, though this does require accurate model predictions of not just the bifurcation points, but also the upcoming changes in the environmental parameters. Between these two extremes in bloom costs, the *reset* strategy can be preferable, allowing a bloom for a limited time, and undoing it later, when conditions are more favourable, saving intervention costs but also ensuring that the bloom does not last all season. Since the *reactive* strategy requires a larger intervention effort then prevention, while also briefly allowing a bloom, it is expected to be the worst option in most cases. Considerable effort could still be saved by returning the control parameter to preventive levels after the lake has switched back to a non-bloom state, but at least the initial effort will always be larger. In practice, however, accurate predictions of the environment are not always available, so prevention is not always an option. In such cases, the reactive strategy could become the best choice. The preventive strategy can also be improved, when considering gradual changes in environmental and control parameters rather than shocks, since the control parameter could be gradually increased to keep pace with the changing environment.

When deciding on a strategy, the stable states are important but do not provide the full picture. As we have shown, the transient dynamics of, e.g., a reset strategy are much faster when the bifurcation point is exceeded by a larger amount. Therefore, it can be favourable to apply a bit more effort than strictly necessary to speed up the recovery, increasing the intervention cost per unit time, but potentially decreasing the overall intervention cost. Simulations of the actual dynamics can play a role in deciding the intensity and duration of a treatment.

To fully realise the potential of our model-based bloom mitigation strategies in practice, two major requirements need to be fulfilled: there must be a model that can accurately predict the bifurcation structure, and there must be a quantitative way to express the costs of interventions and blooms.

In a practical application, the real bifurcation structure will be unknown and the role of the model will be to provide lake managers with an estimate, which must be sufficiently accurate. Of course, achieving this accuracy will require a more detailed model than our example, e.g., PCLake (Janse and van Liere, 1995; Janssen et al., 2019b) and PROTECH (Page et al., 2018), among many others (Rousso et al., 2020; Janssen et al., 2019a). To be used in the way we propose, these models should be calibrated and tested with error measures that reflect their ability to predict bifurcation points, rather than accurate concentrations at given time points, as it is the bifurcation points that are important for the implementation of the strategies (Jacobs et al., 2024). Model uncertainties come from a variety of sources, including the description of the process, the parameter estimates and external variables (Dietze, 2017). Ideally, model uncertainty should be quantified in a way that distinguishes between these different sources, as this allows us to identify where there is room for improvement (Lewis et al., 2022).

Our demonstrations have used a single environmental parameter and a single control parameter. In reality, however, there are many environmental influences acting on a lake (Rousso et al., 2020; Huisman et al., 2018; Burford et al., 2020), and multiple potential control options are available (Lürling and Mucci, 2020). Consequently, the bifurcation structure will be more challenging to understand, analyse and visualise and new tools may be required to deal with, e.g., distances to bifurcation points in multidimensional parameter space.

The other important aspect when choosing a model-based management strategy is translating the costs of blooms and interventions to a numerical value. Not all costs can necessarily be expressed in monetary value. Examples include aesthetic and biodiversity costs to blooms, which different people value to different extents. Estimating these costs therefore involves collecting and combining these different values, a challenging task that can be aided by tools like the Nature Futures Framework (Pereira et al., 2020; Kramer et al., 2023a).

Interventions also have costs that are hard to predict and value. For example, flushing by increasing the flow rate requires diverting upstream water, which can have undesirable effects on water levels or nutrient retention elsewhere (van Wijk et al., 2022). In addition, not all costs are constant over time. For example, diverting a small amount of upstream water for flushing may have no ill effect, while diverting a large amount could be disastrous. Aquatic ecosystem models that connect different lakes can help provide insights into potential flushing costs for the specific lake under consideration (Kramer et al., 2023b).

Finally, the proper design and calibration of a sufficiently accurate model is in itself a difficult and costly task. These costs should also be taken into account, creating a trade-off between a of detailed model-based approach to lake management and more traditional approaches based on expert opinion. This trade-off means that the model-based approach is best applied to lakes with important functions, whether economical (e.g., drinking water, recreation) or ecological (e.g., biodiversity, conservation).

## 5. Conclusion

The presence of alternative stable states with bifurcations in a lake provides interesting opportunities for the short-term model-based mitigation of cyanobacterial blooms, as it allows for qualitatively different strategies. These strategies have different associated cost profiles, so that different strategies can be selected for different situations, depending on their relative costs and the ecological circumstances, which managers could use to their benefit.

## 6. Acknowledgements

We gratefully acknowledge useful discussions with Sven Teurlincx from NIOO-KNAW and Tineke Troost from Deltares. B.J. was funded by project 645.002.002 in the NWO-complexity program.

## Appendix A. Example model equations

The bifurcation diagrams and simulations that we show are from an adapted version of the model for competing green and blue algae by Scheffer et al. (1997a). The model includes competition for light and a single nutrient (phosphorus) of which the total amount *P* is constant and expresses densities of green algae *g* and blue algae *b* in units of nutrient content. We added a low density of both species in the inflow (*g*_*in*_ and *b*_*in*_) to prevent complete extinction and allow switching between stable states. We have also added an environmental parameter σ that reflects the effect of changes in, e.g., temperature or light conditions, as described by Scheffer et al. (1997b). The full equations of the model are:

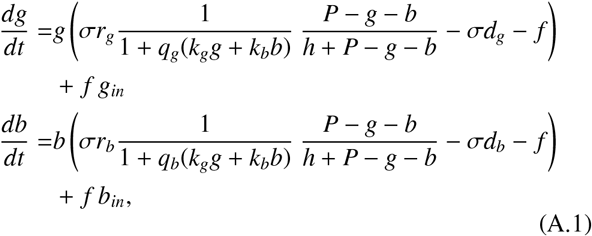

with maximum growth rates *r*_*g*_ and *r*_*b*_, turbidity factors *k*_*g*_ and *k*_*b*_, and sensitivities to turbidity *q*_*g*_ and *q*_*b*_, all for green and blue algae, respectively. The losses due to flow rate (*f*) and half-saturation concentration (*h*) are considered to be the same for both *g* and *b*. For units and default parameter values, see Table A.2.

---

**Table A 2:** Variables and parameters with default values as used in the simulations. N.B., units are taken from Scheffer et al. (1997a), but have no meaning for our demonstration purposes.

| Symbol | Meaning | Value | Unit |
| --- | --- | --- | --- |
| $g(t)$ | Density of green algae | - | $\text{g P} / \text{m}^3$ |
| $b(t)$ | Density of blue algae | - | $\text{g P} / \text{m}^3$ |
| $P$ | Total phosphorus (conserved) | 0.2 | $\text{g P} / \text{m}^3$ |
| $r_g$ | Maximum growth rate of $g$ | 1.2 | $\text{day}^{-1}$ |
| $r_b$ | Maximum growth rate of $b$ | 0.6 | $\text{day}^{-1}$ |
| $h$ | Half-saturation constant | 0.003 | $\text{g P} / \text{m}^3$ |
| $k_g$ | Turbidity factors of $g$ | 5 | $\text{m}^2/\text{g P}$ |
| $k_b$ | Turbidity factors of $b$ | 10 | $\text{m}^2/\text{g P}$ |
| $q_g$ | Sensitivity to turbidity of $g$ | 2 | m |
| $q_b$ | Sensitivity to turbidity of $b$ | 1 | m |
| $d_g$ | Death rate of $g$ | 0.12 | $\text{day}^{-1}$ |
| $d_b$ | Death rate of $b$ | 0.06 | $\text{day}^{-1}$ |
| $f$ | Flow rate | - | $\text{day}^{-1}$ |
| $g_{in}$ | Inflow concentration of $g$ | 0.001 | $\text{g P} / \text{m}^3$ |
| $b_{in}$ | Inflow concentration of $b$ | 0.001 | $\text{g P} / \text{m}^3$ |
| $\sigma$ | Environmental effect | - | - |

